# Interaction of quercetin with transcriptional regulator LasR of Pseudomonas aeruginosa: Mechanistic insights of the inhibition of virulence through quorum sensing

**DOI:** 10.1101/239996

**Authors:** Hovakim Grabski, Lernik Hunanyan, Susanna Tiratsuyan, Hrachik Vardapetyan

## Abstract

**Background:** *Pseudomonas aeruginosa* is one of the most dangerous superbugs in the list of bacteria for which new antibiotics are urgently needed, which was published by World Health Organization. *P. aeruginosa* is an antibiotic-resistant opportunistic human pathogen. It affects patients with AIDS, cystic fibrosis, cancer, burn victims and people with prosthetics and implants. *P. aeruginosa* also forms biofilms. Biofilms increase resistance to antibiotics and host immune responses. Because of biofilms, current therapies are not effective. It is important to find new antibacterial treatment strategies against *P. aeruginosa*. Biofilm formation is regulated through a system called quorum sensing. Thus disrupting this system is considered a promising strategy to combat bacterial pathogenicity. It is known that quercetin inhibits *Pseudomonas aeruginosa* biofilm formation, but the mechanism of action is unknown. In the present study, we tried to analyse the mode of interactions of LasR with quercetin.

**Results:** We used a combination of molecular docking, molecular dynamics (MD) simulations and machine learning techniques for the study of the interaction of the LasR protein of *P. aeruginosa* with quercetin. We assessed the conformational changes of the interaction and analysed the molecular details of the binding of quercetin with LasR. We show that quercetin has two binding modes. One binding mode is the interaction with ligand binding domain, this interaction is not competitive and it has also been shown experimentally. The second binding mode is the interaction with the bridge, it involves conservative amino acid interactions from LBD, SLR, and DBD and it is also not competitive. Experimental studies show hydroxyl group of ring A is necessary for inhibitory activity, in our model the hydroxyl group interacts with Leu177 during the second binding mode. This could explain the molecular mechanism of how quercetin inhibits LasR protein.

**Conclusions:** This study may offer insights on how quercetin inhibits quorum sensing circuitry by interacting with transcriptional regulator LasR. The capability of having two binding modes may explain why quercetin is effective at inhibiting biofilm formation and virulence gene expression.

**List of abbreviations:**

PDB: Protein data bank
MD: Molecular Dynamics
PCA: Principal Component Analysis
PC: Principal Component
SLR: Short Linker Region
BLAST: Basic local alignment search tool
DBI: David-Bouldin Index
psF: pseudo-F statistic

## Background

*Pseudomonas aeruginosa* is one of the “ESKAPE” pathogens and has acquired resistance to commonly used antibiotics [1, 2]. It is vital to find ways on how to counteract against it. *P. aeruginosa* is an opportunistic human pathogen and of clinical relevance, because it affects people with cystic fibrosis, cancer, burn victims, with implants and prosthetics, etc. *P. aeruginosa* uses quorum sensing system for the regulation of collective behaviors. This system controls virulence factor production. *P. aeruginosa* is pathogenic because of the synthesis of virulence factors such as proteases, rhamnolipids, hemolysins, production of antibiotic pyocyanin, Hydrogen Cyanide (HCN), secretion systems of Types 1 (T1SS), 2 (T2SS), 3 (T3SS), 4[3], 5 (T5SS), and 6 (T6SS) [4], and biofilm formation [3,5]. *P. aeruginosa* QS circuit includes transcriptional regulator LasR and RhIR, which detect 3-O-C12 homoserine lactone and C4 homoserine lactone [6–9]. In our previous research, we show that there are multiple binding modes of the native ligand with the transcriptional activator LasR [10].

There have been numerous attempts for the development of *P. aeruginosa* QS inhibitors [11–14]. These efforts resulted in the findings of inhibitors that work in vitro, but not in vivo models in animals [15]. Most of these researches assume that the inhibitors bind to the ligand binding domain (LBD). There was also another research involving flavonoids as inhibitors of biofilm formation [16]. Flavonoids are a group of natural products and secondary metabolites that exhibit broad spectrum of pharmacological activities such as antimicrobial, antiinflammatory, etc [17]. However, their mechanism of action is not well investigated. One such flavonoid is quercetin, which is considered as generally safe compound [16].

Hence, we analysed the molecular details of the interactions of quercetin with LasR protein. So far this is the first report that shows that the quercetin can interact as well with the bridge of LasR [10]. This study may explain why and how quercetin inhibits biofilm formation on a molecular level.

## Methods

### LasR and quercetin models

Previously modeled LasR monomer structure was used for study [10]. The 2D and 3D structure of quercetin were obtained from PubChem [18] and they are shown in Figure 1.

**Fig. 1.**
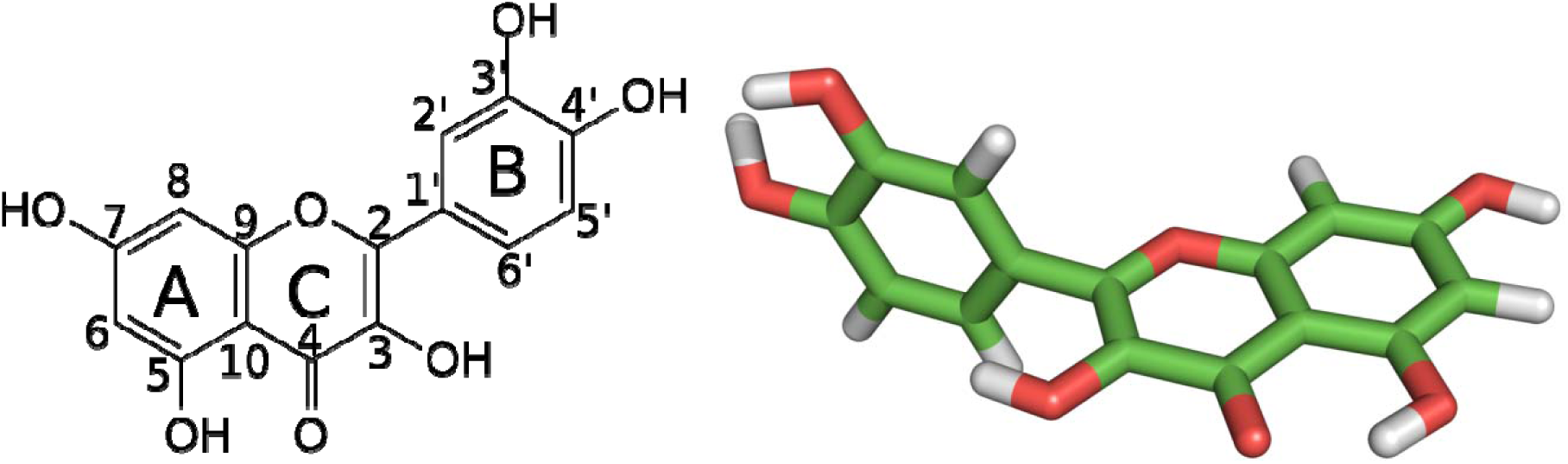
Quercetin molecule structure: left) 2D right) 3D stick representation.

The ligand parameters were generated using the acpype tool [19] for the General Amber Force Field [20] with AM1-BCC partial charges [21].

### LasR–quercetin ligand blind docking experiments

Docking experiments of quercetin with LasR monomer were carried out using Autodock Vina [22, 23], and it is based on the use of a rigid receptor. The whole protein conformational space was searched, using grid box dimensions 60×62×48 A°. Following exhaustiveness values were tested in this study: 8, 16, 32, 64, 128, 256, 512, 1024, 2048 and 4096. Principal component (PC) [24] and cluster analysis, using K-means algorithm [25], were performed (Additional file 1: Figure S1). The number of interaction sites doesn’t change in the interval using exhaustiveness from 1024 to 4096. Exhaustiveness value of 1024 was chosen as it provides good results, good speed and thorough sampling of the docked configurations.

Exhaustiveness value was increased to a value of 1024, and a maximum number of binding modes to generate set to 20. After that 100 independent docking calculations were carried out with random initial seeds. Later we verified blind docking results with rDock [26] and FlexAid [27].

### Molecular dynamics simulations of LasR–quercetin systems

The same methodology was used from our previous work [10]. We conducted the MD simulations with the GROMACS suite, version 5.1.2 [28] Amber ff99SB-ILDN force field [29] was used for the MD simulations. In all cases, Short-range non-bonded interactions were cut off at 1.4 nm. Particle Mesh Ewald (PME) [30, 31] was used for the calculation of long-range electrostatics. A time step of 2 fs was used during heating, the production run, and coordinates were recorded every 1 ps. Two simulations of 300 ns were performed.

Structures were placed in a dodecahedron box of TIP3P water [32], to which 100 mM NaCl was added, including neutralizing counter-ions. After that two steepest descents minimization were performed and then equilibrated in two stages. The first stage involved simulating for 200 ps under a constant volume (NVT) ensemble. The second stage involved simulating for 200 ps under a constant-pressure (NPT) for maintaining pressure isotropically at 1.0 bar. The temperature was sustained at 300 K using V-rescale [33] algorithm. For isotropic regulation of the pressure, the Parrinello-Rahman barostat [34] was used.

### Sequence conservation

To find out whether quercetin interacts with conservative amino acid residues we performed multiple sequence alignment (MSA) using msa package [35]. ClustalW [36], Clustal Omega [37] and Muscle [38] within msa package were used for multiple sequence alignments. We performed multiple sequence alignment of the LasR protein between the closely related species of *P.aeruginosa*.

The following sequences were used for sequence alignment:

- WP_054058449.1 LuxR family transcriptional regulator [*Pseudomonas fuscovaginae*]
- NP_250121 transcriptional regulator LasR [*Pseudomonas aeruginosa* PAO1]
- KFC75736.1 LuxR-type transcriptional regulator [Massilia sp. LC238]
- WP_018433960 LuxR family transcriptional regulator [Burkholderia sp. JPY251]
- WP_042326260.1 LuxR family transcriptional regulator [Paraburkholderia ginsengisoli]
- WP_012426170.1 LuxR family transcriptional regulator [Paraburkholderia phytofirmans]
- WP_027776298.1 LuxR family transcriptional regulator [Paraburkholderia caledonica]
- WP_027214716.1 LuxR family transcriptional regulator [Burkholderia sp. WSM2232]
- WP_041729325.1 LuxR family transcriptional regulator [Burkholderia sp. CCGE1003]
- CBI71275.1 UnaR protein [Paraburkholderia unamae]
- CAP91064.1 BraR protein [Paraburkholderia kururiensis]
- WP_003082999.1 transcriptional activator protein LasR [*Pseudomonas*]
- WP_012076422.1 MULTISPECIES: LuxR family transcriptional regulator [*Pseudomonas*]
- WP_050376898.1 MULTISPECIES: LuxR family transcriptional regulator [*Pseudomonas*]
- WP_050395760.1 LuxR family transcriptional regulator [*Pseudomonas aeruginosa*]

### Entire flowchart

The whole methodology is based on our previous research [10] and is presented as a flowchart for a better comprehension:

- The reconstructed model was from our previous research.
- The 3D model of quercetin acquired from pubchem web server.
- Blind docking of quercetin with the LasR monomer performed using Autodock Vina.
- Later verified blind docking with rDock and FlexAid.
- PCA and cluster analysis performed on docking conformations.
- Extraction of centroid conformations from cluster analysis.
- Ligand parameters generated using acpype interface in the framework of the AMBER force field.
- Using centroid conformations as starting points for molecular dynamics simulations using Gromacs.
- Analysis of molecular dynamics trajectory files using MDTraj.
- Binding energy calculated using g_mmpbsa
- Sequence conservation analysis performed using msa library.

## Results and discussion

### Docking analysis of quercetin with LasR monomer

Molecular docking was used for the prediction of binding modes of quercetin with LasR monomer. PCA and cluster analysis were performed on docking data using center coordinates of the conformations (Additional file 1: Figure S2). There are four binding sites, cluster 1 and 3 correspond to the interaction with LBD [16], cluster 2 corresponds to the interaction with LBD-SLR-DBD bridge [10], cluster 4 is not significant since it only encompasses only 0.5% of the docking simulations.

Several rounds of K-means clustering (details are available in the section of methods) were performed using the docking data. The accuracy of the cluster analysis was evaluated using the DBI [39], Dunn Index [40], Silhouette score [41] and the pSF [42] metrics (Additional file 1: Figure S3). An optimal number of clusters were chosen for docking results, simultaneously accounting for a low DBI, high Dunn, high Silhouette and high pSF values.

We generated 1979 docked poses and performed representative structure extraction for use in MD simulations of the LasR–quercetin binding sites. The resulting representative structures from each cluster are shown in Figure 2. These cluster representative structures were produced by finding the centroid conformations.

**Fig. 2.**
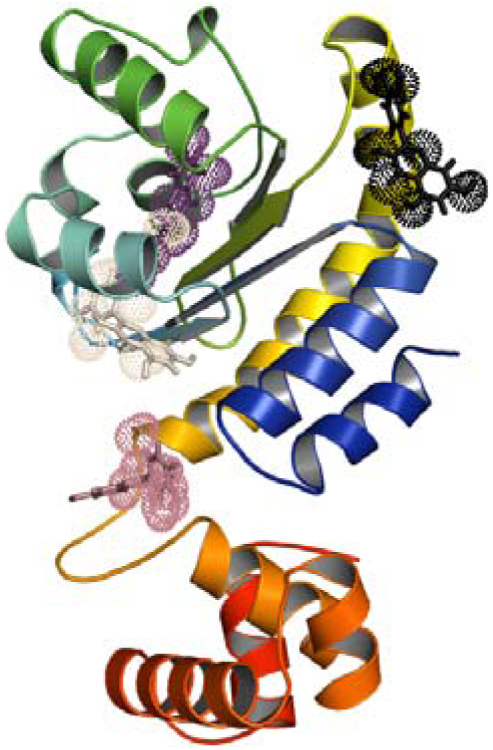
3D visualization of the analysed docking data with their representative structures and clusters.

Representative structures from each cluster were extracted. The binding energy for the representative structure of cluster 1 is -7.6 kcal/mol, as for the mean binding affinity for the whole cluster is -7.206 (SD 0.398) kcal/mol (Additional file 1: Figure S4). Cluster 1 contains 946 docked poses from 1979, about 47.802%. For cluster 2 the binding affinity for the representative structure is -6.8 kcal/mol and for the whole cluster -6.971 (SD 0.334) kcal/mol (Additional file 1: Figure S5). Cluster 2 contains 920 docked poses from 1979, about 46.488%, which is a rather unstudied area. For cluster 3, the representative structure features the highest binding affinity - 7.8 kcal/mol and for the whole cluster -7.793 (SD 0.072) kcal/mol (Additional file 1: Figure S6). Cluster 3 contains 100 docked poses from 1979, about 5.053%. For cluster 4, the representative structure features the highest binding affinity -6.6 kcal/mol and for the whole cluster -6.592 (SD 0.027) kcal/mol (Additional file 1: Figure S7). Cluster 4 contains 13 docked poses from 1979, about 0.657%.

Unfortunately molecular docking is not appropriate for the prediction of binding affinity or binding poses of protein-ligand complexes, however, they can still provide important information [10, 43–44]

For molecular dynamics simulations, we only used centroid conformations from cluster 1 and 2, since they encompass 94.29% of the docking data.

Later we corroborated the blind docking with other molecular docking software as well, which includes Autodock Vina [22], rDock [26] and FlexAid [27] (Figure 3). PC analysis of the various docking programs was performed for easier comprehension (Additional file 1: Figure S8).

**Fig. 3.**
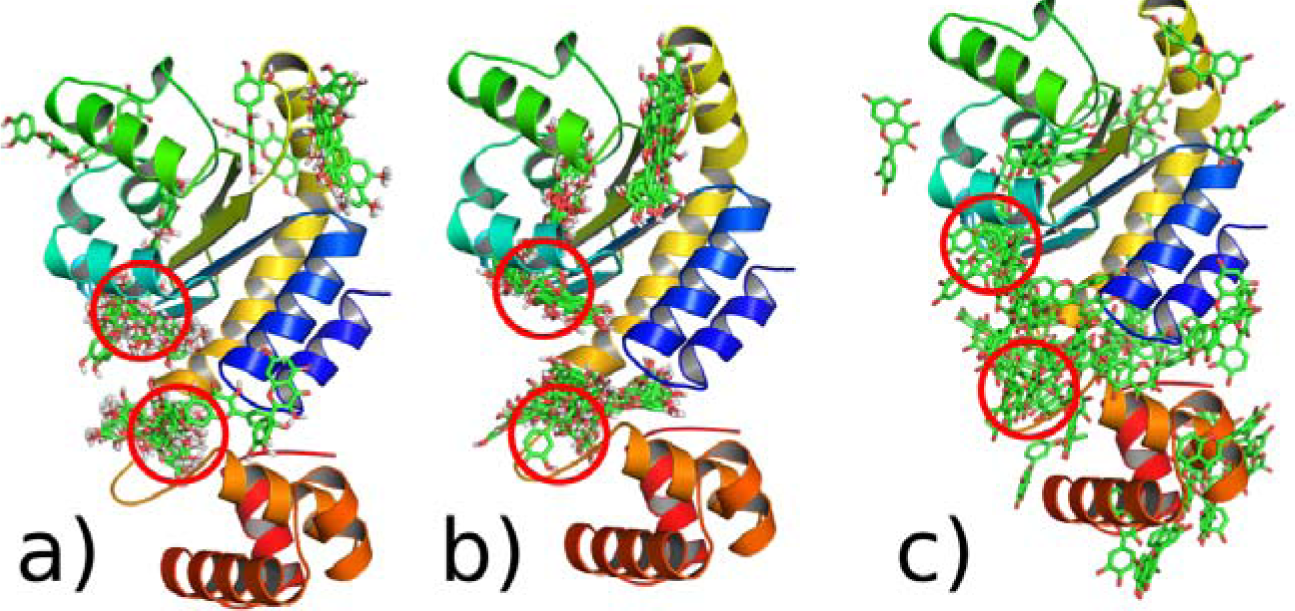
3D visualization of the analysed docking data with their representative structures and clusters.

### Binding modes of quercetin

We performed two 300 ns simulations using standard MD protocol. Overall, 600ns of aggregate simulation data was used for the analysis of the interaction quercetin with LasR monomer. The representative structures from docking results (Figure 2) were used as starting points for MD simulation with LasR. We used the mass-weighted RMSD for the assessment of the overall stability of the ligand. RMSD was calculated with reference to the initial snapshot for the different independent MD runs.

Principal component analysis was performed using molecular dynamics conformations. We used the cumulative proportion to assess the total amount of variance that the consecutive principal components explain. After that several rounds of agglomerative clustering (details are available in the section of methods) were performed using the simulation data. The accuracy of the cluster analysis was evaluated using the DBI [39], Dunn Index [40], Silhouette score [41] and the pSF [42] metrics. An optimal number of clusters was chosen for molecular dynamics data sets, simultaneously accounting for a low DBI, high Dunn, high Silhouette and high pSF values.

***Interaction of quercetin with LBD***. The first two principal components explain 93.34% of the cumulative proportion variance for the conformational changes of quercetin (Additional File 1: Figure S9). Histogram analysis of the RMSD evolution shows there are two peaks (Additional File 1: Figure S10), this is another confirmation for cluster analysis (Additional File 1: Figure S11-S12).

Hydrogen and hydrophobic analysis show that nine amino acid residues interact with the ligand. Three amino acid residues, which include Arg61, Ala50, Glu48, form hydrogen bonds. Six amino acid residues interact hydrophobically with quercetin (Figure 4).

**Fig. 4.**
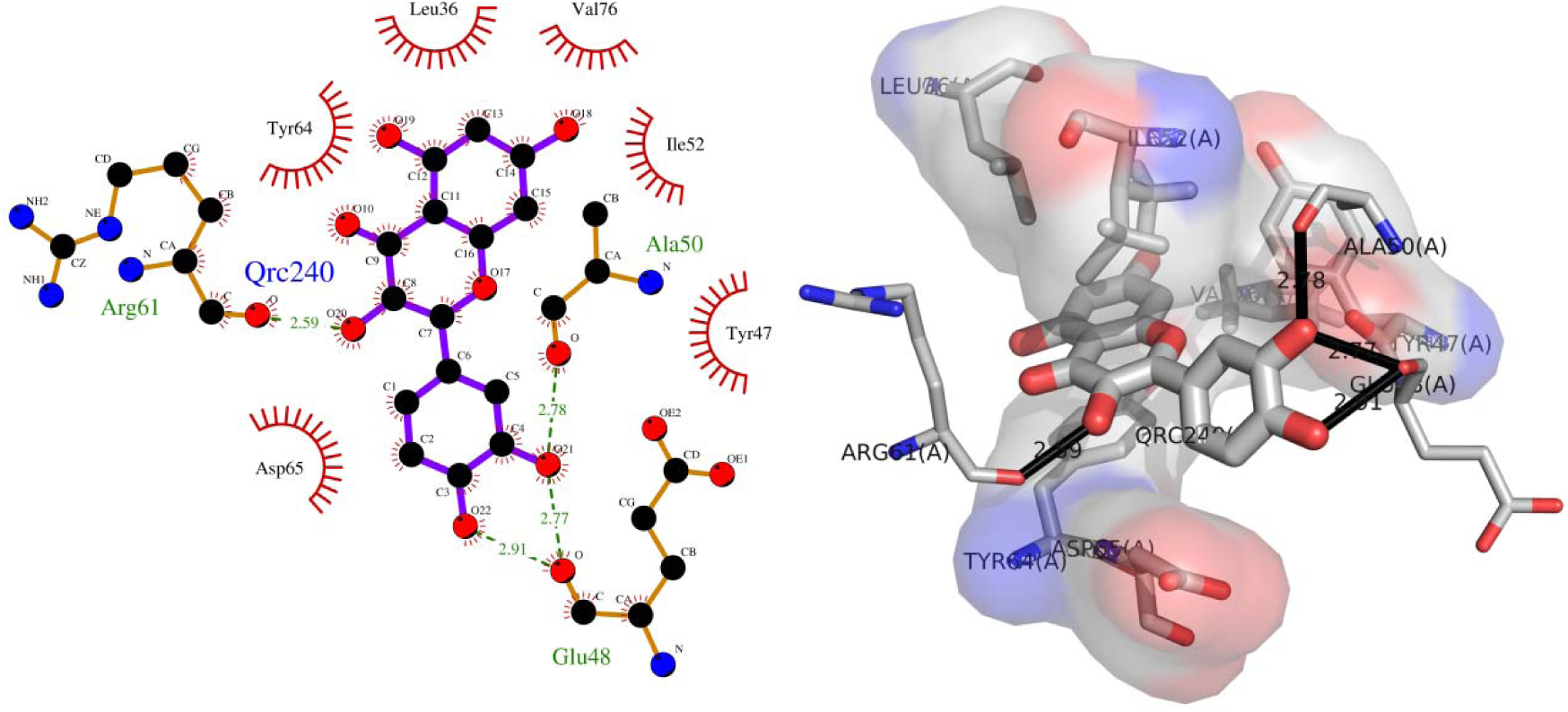
Schematic and 3D representation of hydrogen and hydrophobic interactions of quercetin with LBD of LasR.

Quercetin does not enter the binding pocket fully (Figure 5), this could suggest for a ternary interaction possibility, so has been shown experimentally [16].

**Fig. 5.**
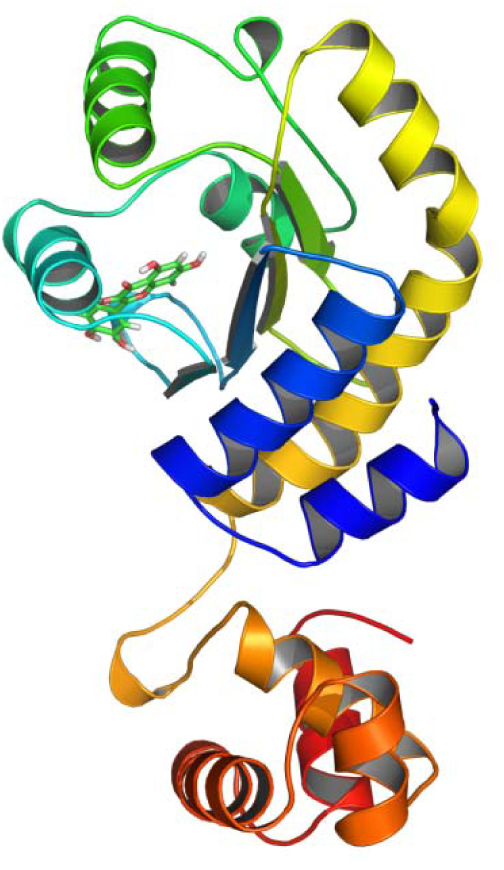
Quercetin interaction with LBD of LasR.

***Interaction of quercetin with the bridge***. The first two principal components explain 90.7% of the cumulative variance for the conformational changes of quercetin (Additional File 1: Figure S13). Histogram analysis of the RMSD evolution shows there are two peaks (Additional File 1: Figure S14), this is another confirmation for cluster analysis (Additional File 1: Figure S15-S16).

Hydrogen and hydrophobic analysis show that thirteen amino acid residues interact with the ligand. Three amino acid residues, which include Glu11, Ser13, Leu177, form hydrogen bonds. Ten amino acid residues interact hydrophobically with quercetin (Figure 6).

**Fig. 6.**
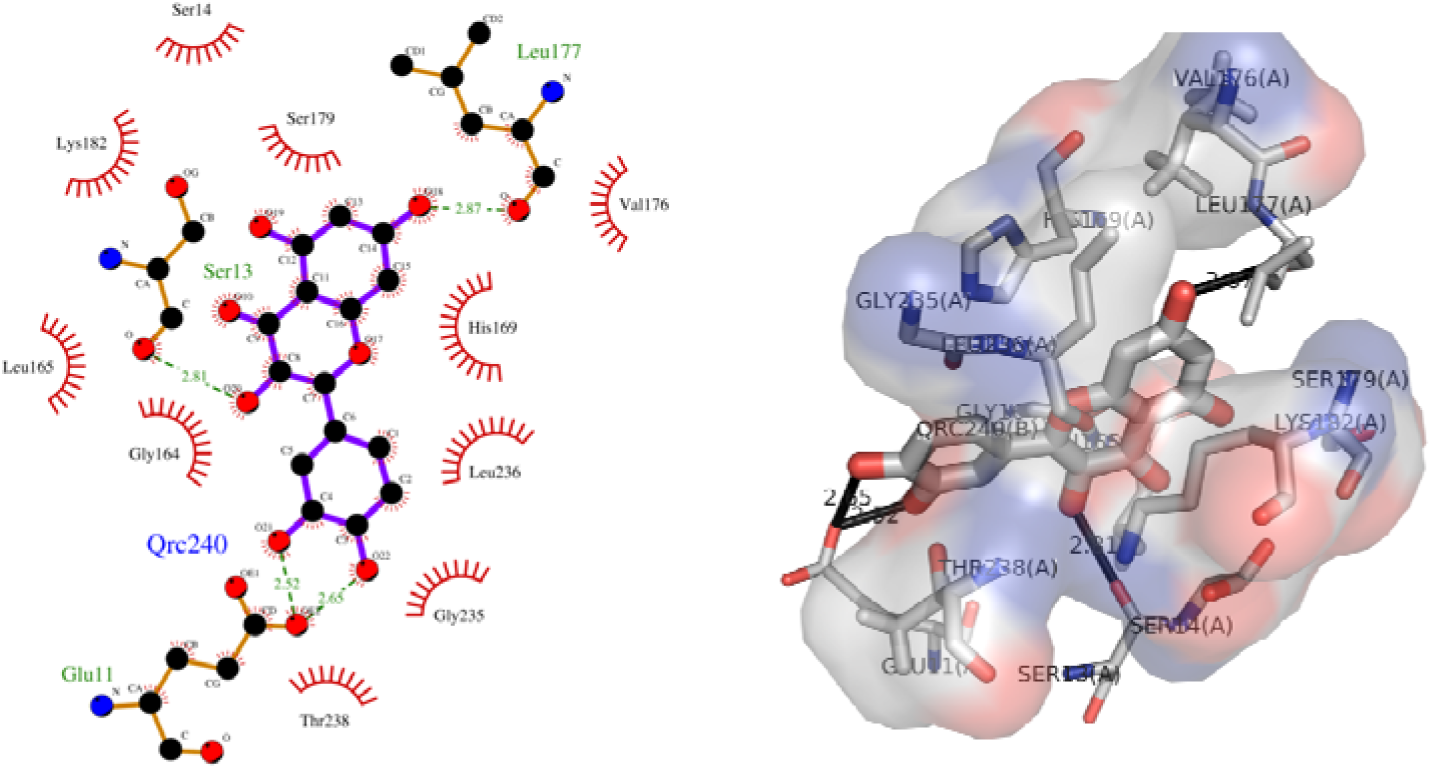
Schematic and 3D representation of hydrogen and hydrophobic interactions of quercetin with the bridge of LasR.

Quercetin interacts with the bridge, just like 3-O-C12 HSL [10] (Figure 7).

**Fig. 7.**
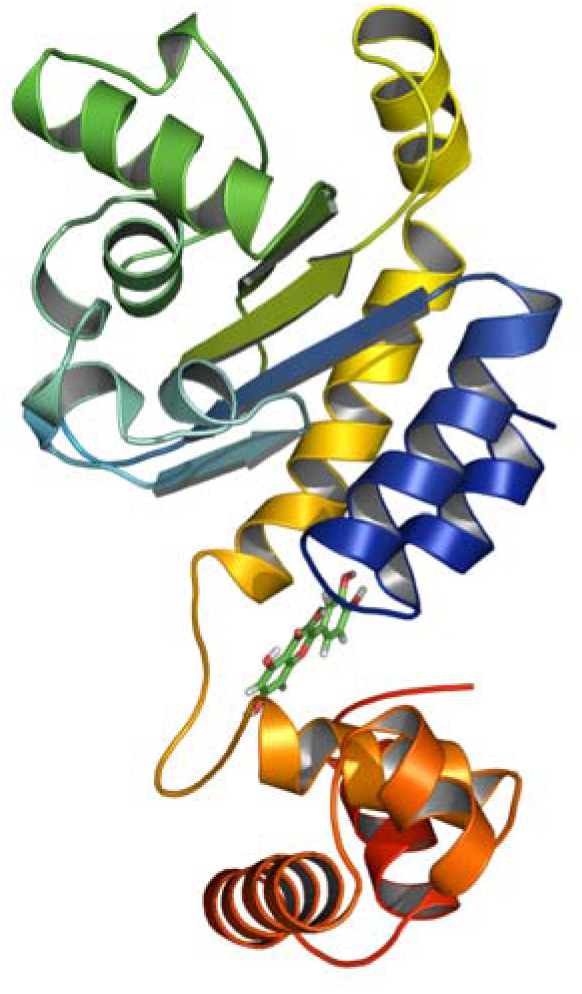
Interaction of quercetin with the bridge of LasR.

### Binding energy of quercetin to LasR and sequence conservation

In order to analyse the binding sites in detail, MMPBSA [45] binding energy calculation was performed for each binding site based on the trajectories. It is also interesting that quercetin does not compete for the LBD (Table 1), which has also been shown experimentally [16]. But the results suggest that the interaction of quercetin with the bridge is not competitive, but allosteric.

More detailed analysis of the energy terms showed that Van der Waals, electrostatic interactions, and non-polar solvation energy contribute negatively to the binding energy while polar solvation energy contributes positively. Electrostatic interaction contributes most in the terms of negative contribution for both cases, but for the interaction with the LBD-SLR-DBD bridge or “the bridge”, the electrostatic interaction is 1.95 times higher than with the LBD interaction.

**Table 1.** Relative Binding energy using g_mmpbsa on simulation data.

| Binding sites | Van der waals<br>(kJ/mol) | Electrostatic<br>(kJ/mol) | Polar<br>salvation<br>(kJ/mol) | Non-polar<br>salvation<br>(kJ/mol) | Binding energy<br>(kJ/mol) |
| --- | --- | --- | --- | --- | --- |
| 3-O-C12 HSL-LBD | -215.673±8.007 | -39.586±18.817 | 147.522±13.978 | -20.840±0.794 | -128.578±18.757 |
| 3-O-C12 HSL-bridge | -205.141±11.803 | -142.974±29.032 | 201.889±20.117 | -20.230±0.842 | -166.456±20.492 |
| QRC-LBD | -148.890±14.038 | -123.004±33.612 | 158.564±14.347 | -16.113±0.657 | -129.443±19.319 |
| QRC-bridge | -129.937±15.771 | -240.057±23.884 | 201.062±11.982 | -14.983±0.545 | -183.915±16.249 |

We performed analysis of energy contribution of residues to binding for both simulations obtained from MMPBSA calculation using g_mmpbsa [45]. Sequence alignment was performed using msa package [35]. ClustalW [36], Clustal Omega [37] and Muscle[38] algorithms were used for sequence alignment. The binding state with LBD with binding energy ~ -129.443 kJ/mol (Table 1), which the residues Asp65, Tyr64, ILE52, GLU48, and Arg61 contribute most to (Figure 8).

**Fig. 8.**
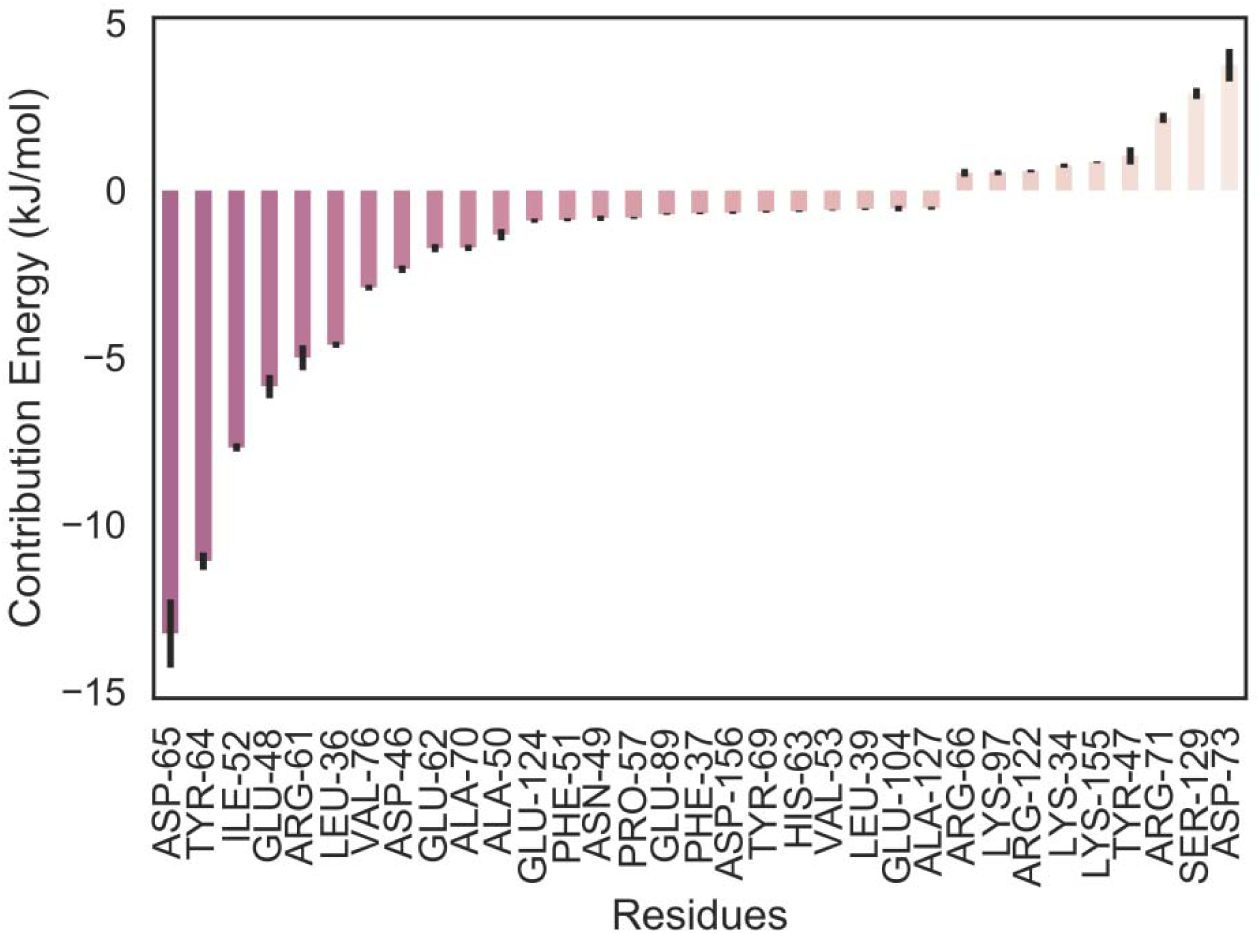
Energy contribution of LBD amino acid residues from simulation 1.

Residue interaction and sequence alignment show that quercetin interacts with very conservative amino acid that includes Tyr64, Arg61, Asp73, and others (Figure 9). Quercetin interacts with 7 fully conserved amino acids. In total 10 amino acids participate in the interaction, where amino acid conservation is more than 75%.

**Fig. 9.**
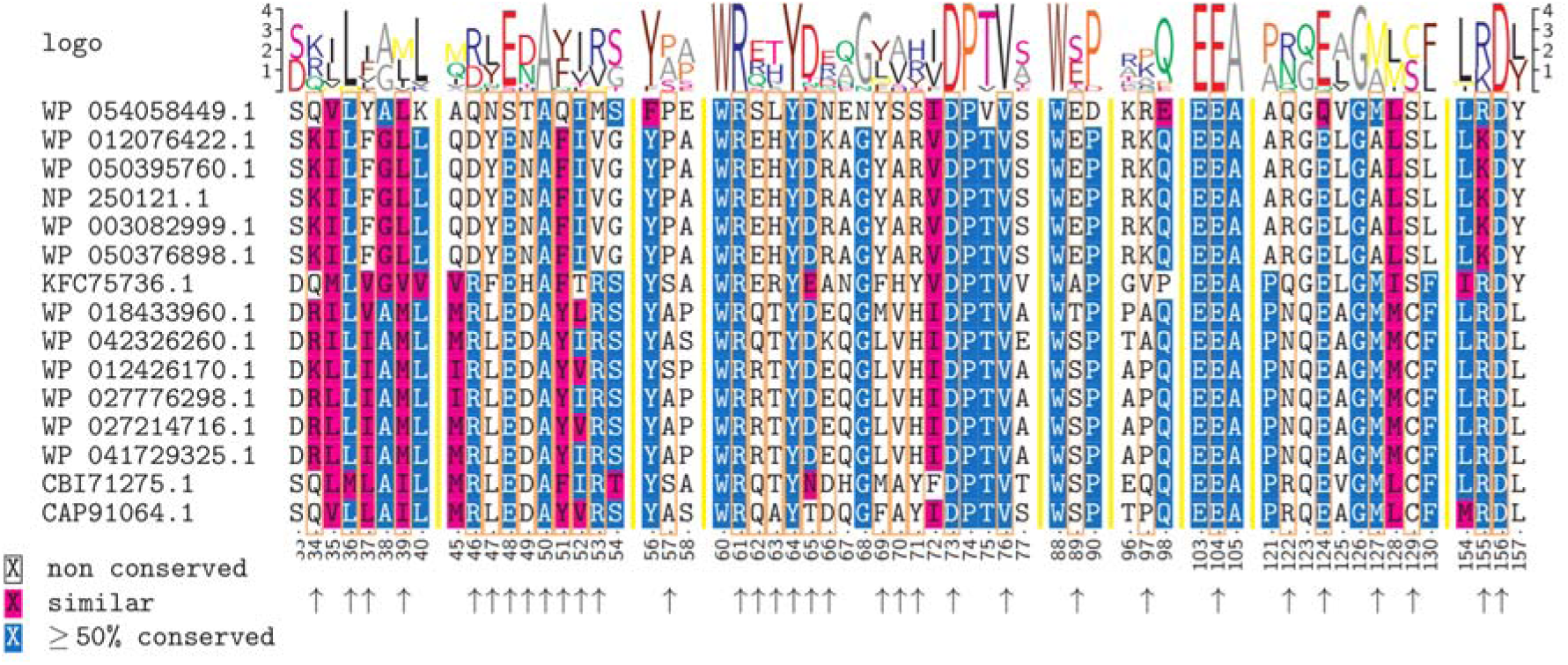
Interaction of conserved amino acids of LBD of LasR with quercetin molecule. Boxes and arrows point the amino acid residues that interact with quercetin.

Then we performed analysis of which rings of the quercetin molecule participate in the interaction with the amino acids. Aps65 interacts with hydroxyl group of ring C at position 3 and hydroxyl groups of ring B at positions 3’ and 2’ (Figure 1). Tyr64 interacts with atoms from ring C: with hydroxyl groups at position 3 and with oxygen at position 1 and 4. Tyr64 also interacts with ring A: with hydroxyl groups at position 5. Ile52 interacts with ring C and B: with hydroxyl group at position 3 of ring C, and hydrogen at position 6’ of ring B. Arg61 interacts with ring C: oxygen at position 4 and hydroxyl group at position 3. Leu36 interacts with ring A: with hydrogen at position 6 and hydroxyl group at position 5. Val76 which is a conservative residue interacts with ring A: with hydroxyl group at position 7. Amino acids that contribute positively include Asp73 and Ser129. Asp73, which is conservative residue, interacts with ring A: with hydroxyl group at position 7. The hydroxyl group at position 7 is necessary for inhibitory activity and has been shown experimentally [16].

The new binding state with LBD-SLR-DBD bridge or “the bridge” of LasR with energy ~ -183.915 kJ/mol (Table 1), where residues Glu11, Leu177, Leu236, Lys182, Lys218 contribute most (Figure 10).

**Fig. 10.**
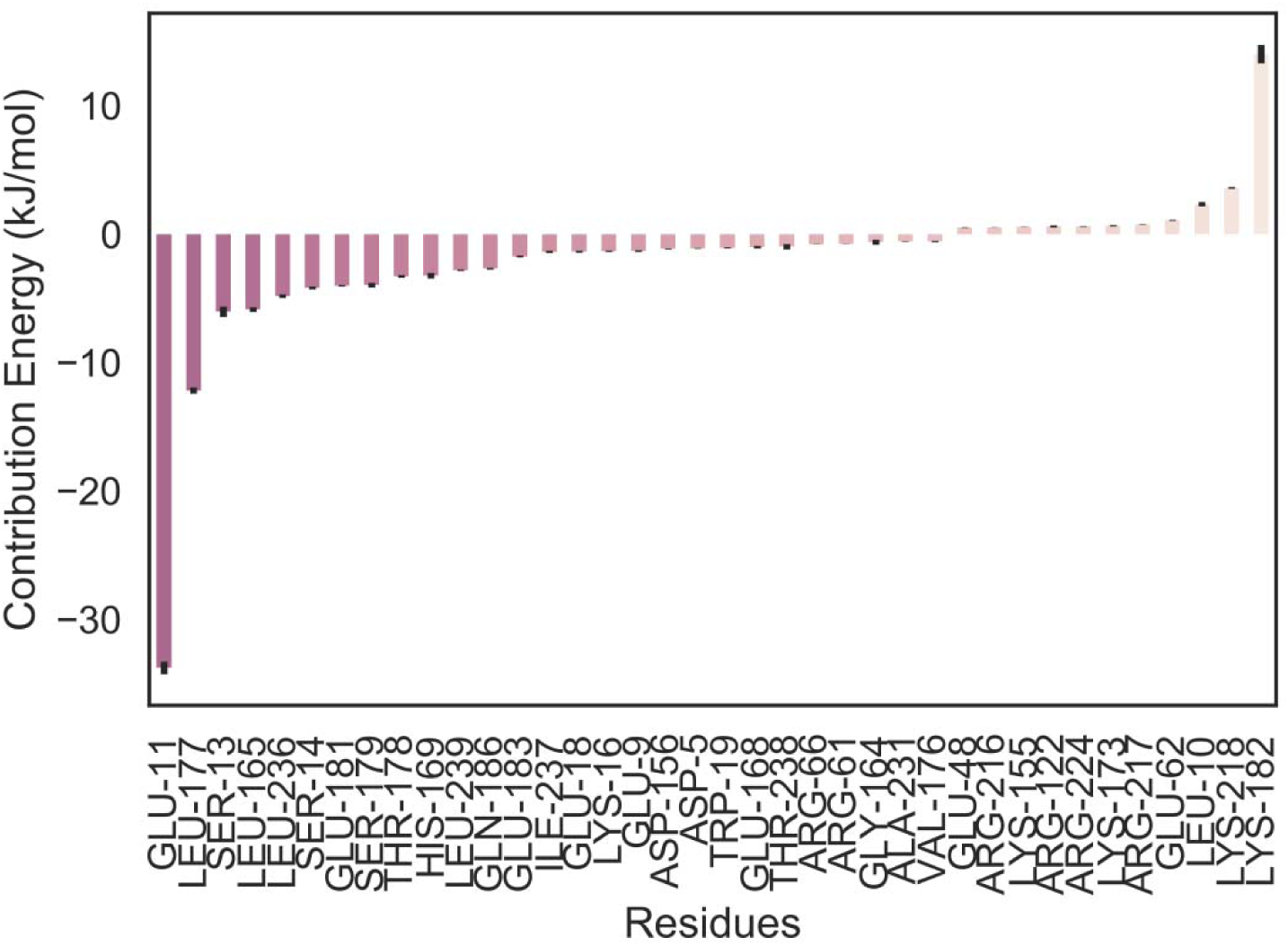
Energy contribution of LBD-SLR-DBD bridge amino acid residues from simulation 2.

Analysis of residue sequence alignment show quercetin interacts with very conservative amino acids such as Leu236, Leu177, Phe219, Trp19 (Figure 11). Quercetin interacts with 12 fully conserved amino acids. In total 16 amino acids participate in the interaction, where amino acid conservation is more than 75%.

**Fig. 11.**
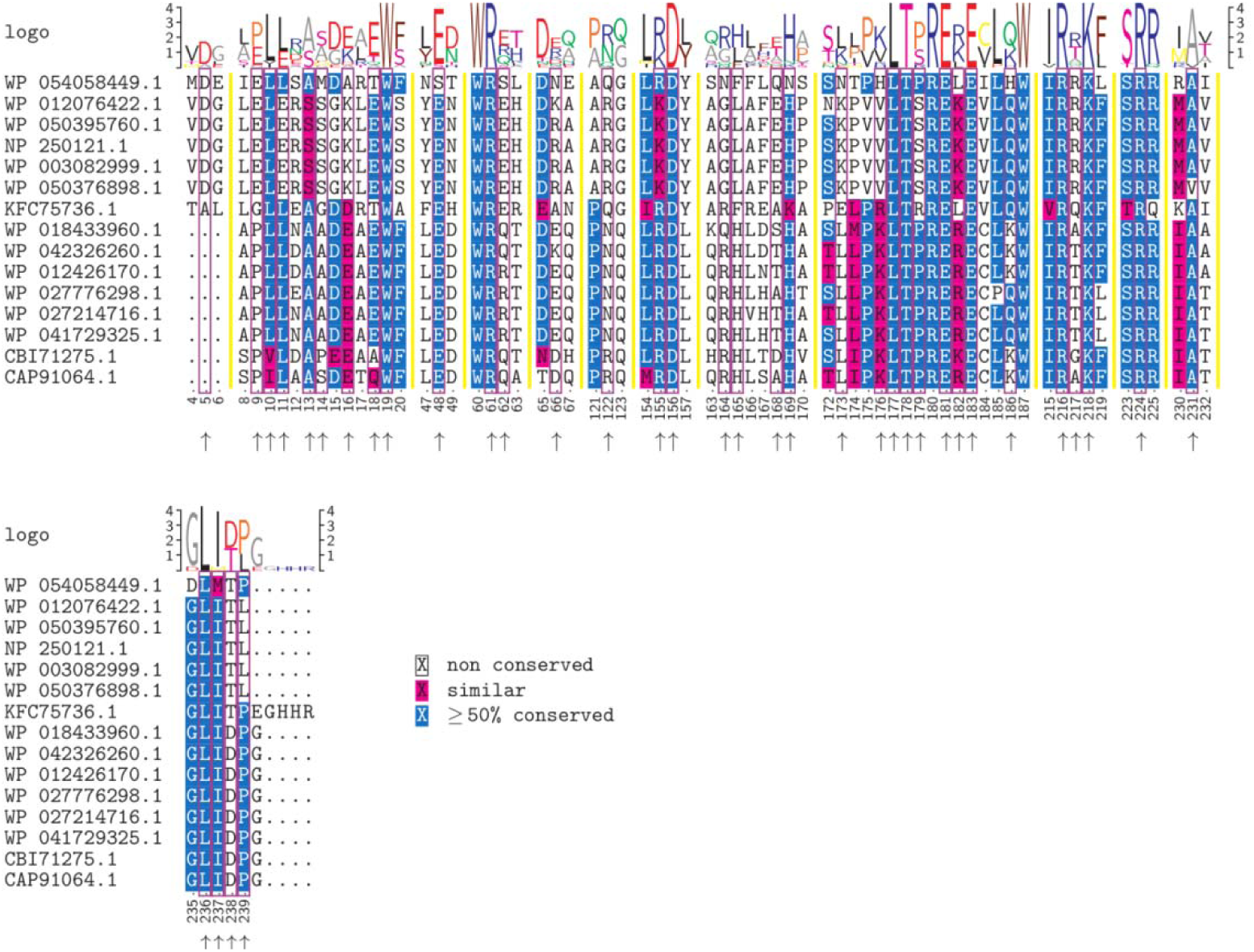
Interaction of conserved amino acids of LBD-SLR-DBD bridge (“the bridge”) with the autoinducer molecule. Boxes and arrows point the residues that interact with quercetin.

In this new binding site Glu11 from LBD, Leu177 from SLR and Lys236 of DBD participate in the interactions with LasR. Then we performed analysis of which parts of the quercetin molecule interact with the amino acid residues. Glu11 interacts with ring B: mainly with carbon atoms and hydroxyl groups at position 4’ and 3’. Leu177 interacts with ring A: especially carbon atoms at position 7 and 6 and hydroxyl group at position 7. Leu236 interacts with rings A, B, and C: carbon and hydroxyl groups at position 7 of ring A, hydrogen of ring B at position 6’, carbon and hydrogen at position 8 of ring C. Lys182 interacts with ring A and C: carbon atoms of ring A at position 7 and 8. This result clearly suggests that both of the C-terminal and the N-terminal of LasR interact with quercetin.

## Conclusion

From the simulations, it can be safely concluded that quercetin can bind both to LBD and to “bridge” of transcriptional regulator LasR. This suggests that there are multiple binding modes rather than one. From experimental studies, it has been shown that hydroxyl group at position 7 of ring A is important for inhibitory activity. In our case, it is visible that quercetin interacts with Leu177 with hydroxyl group at position 7 from ring A. This amino acid residue is a conservative and from the short linker region between LDB and DBD. This could explain how quercetin inhibits DNA binding by preventing hinge rotation of DBD.

The interaction with the LBD-SLR-DBD bridge is a novel site. The analysis of binding energy shows that the interaction of quercetin with “bridge” is not competitive. Conservative amino acids such as Leu177, Leu236, Lys182, Lys218 contribute most during the interaction with LBD-SLR-DBD bridge. This could suggest that for the inhibition of DNA binding capability, it is necessary the interaction of quercetin with the “bridge”.

This study may reveal new insights of the interactions of the quercetin with transcriptional regulator LasR of *P. aeruginosa*. Results from this study may explain why quercetin is effective at inhibiting transcriptional regulator LasR and stop biofilm and virulence gene expression.

## Additional files

**Additional file 1: Figure S1**. Determination of exhaustiveness after PCA and cluster analysis. **Figure S2**. Clustering results on the docking data using k-means algorithm formed by first two PCs. **Figure S3**. K-means clustering with a different number of clusters for the quercetin docking data. **Figure S4**. Mean Autodock Vina binding energy score for Cluster 1 and percentage from total of conformations **Figure S5**. Mean Autodock Vina binding energy score for Cluster 2 and percentage from total of conformations. **Figure S6**. Mean Autodock Vina binding energy score for Cluster 3 and percentage from total of conformations. **Figure S7**. Mean Autodock Vina binding energy score for cluster 4 and percentage from the total of conformations. **Figure S8**. Principal component analysis of ligand (quercetin) conformations obtained from various docking programs. **Figure S9**. Cumulative proportion of the components for quercetin conformations during the interaction with LBD of LasR. **Figure S10**. RMSD histogram of quercetin during the interaction with LBD. **Figure S11**. PCA and RMSD evolution of quercetin during 300 ns of simulation during the interaction with LBD using representative structure of quercetin docking pose as starting point. **Figure S12**. Quality of agglomerative clustering with a different number of clusters for the quercetin-LBD MD data. **Figure S13**. Cumulative proportion of the components for quercetin conformations during the interaction with the bridge of LasR. **Figure S14**. RMSD histogram of quercetin during the interaction with the bridge. **Figure S15**. PCA and RMSD evolution of quercetin during 300 ns of simulation during the interaction with the bridge using representative structure of quercetin docking pose as starting point. **Figure S16**. Quality of agglomerative clustering with a different number of clusters for the quercetin-bridge MD data. (DOCX)

**Additional file 2:** Movie of MD simulation 1. (MP4)

**Additional file 3:** Movie of MD simulation 2. (MP4)

## Declarations

### Acknowledgments

We thank Yerevan Physics Institute for providing time on the cluster for the molecular dynamics simulations.

### Availability of data and materials

The analysed datasets are available as additional files.

